# Dityrosine cross-links are present in Alzheimer’s disease-derived tau oligomers and paired helical filaments (PHF) which promotes the stability of the PHF-core tau (297-391) *in vitro*

**DOI:** 10.1101/2022.05.28.493839

**Authors:** Mahmoud B. Maina, Youssra K. Al-Hilaly, Sebastian Oakley, Gunashekar Burra, Tahmida Khanon, Luca Biasetti, Kurtis Mengham, Karen Marshall, Janet E. Rickard, Charles R. Harrington, Claude M. Wischik, Louise C. Serpell

## Abstract

A characteristic hallmark of Alzheimer’s Disease (AD) is the pathological aggregation and deposition of tau into paired helical filaments (PHF) in neurofibrillary tangles (NFTs). Oxidative stress is an early event during AD pathogenesis and is associated with tau-mediated AD pathology. Oxidative environments can result in the formation of covalent dityrosine crosslinks that can increase protein stability and insolubility. Dityrosine cross-linking has been shown to occur *in vivo* in Aβ plaques and α-synuclein aggregates in Lewy bodies, and this modification may increase the insolubility of these aggregates and their resistance to degradation. Using the PHF-core tau fragment (residues 297 – 391) as a model, we have previously demonstrated that dityrosine formation traps tau assemblies to reduce further elongation. However, it is unknown whether dityrosine crosslinks are found in tau deposits *in vivo* in AD and its relevance to disease mechanism is unclear. Here, using transmission electron microscope (TEM) double immunogold-labelling, we reveal that neurofibrillary NFTs in AD are heavily decorated with dityrosine crosslinks alongside tau. Single immunogold-labelling TEM and fluorescence spectroscopy revealed the presence of dityrosine on AD brain-derived tau oligomers and fibrils. Using the tau (297-391) PHF-core fragment as a model, we further showed that prefibrillar tau species are more amenable to dityrosine crosslinking than tau fibrils. Dityrosine formation results in heat and SDS stability of oxidised prefibrillar and fibrillar tau assemblies. This finding has implications for understanding the mechanism governing the insolubility and toxicity of tau assemblies *in vivo*.

## Introduction

Tau protein was first described as a cytoplasmic protein that plays a role in microtubule dynamics (Cleveland et al., 1977, Weingarten et al., 1975). However, recent evidence suggests that it plays multiple functions, including DNA protection, heterochromatin stability and nucleolar transcription (Maina et al., 2018, Sotiropoulos et al., 2017, Bukar Maina et al., 2016, Sultan et al., 2011). Interest in tau intensified following the discovery that it is the main component of the intracellular neurofibrillary tangles (NFTs), comprised of straight (SFs) and paired-helical filaments (PHFs), which forms one of the hallmarks of Alzheimer’s disease (AD), alongside extracellular deposits of amyloid-beta (Aβ) (Grundke-Iqbal et al., 1986, Wischik et al., 1988, Brion et al., 1985, Kosik et al., 1986). Under normal conditions, tau is soluble and intrinsically disordered. However, post-translational modifications, truncation in disease, mutations or cellular stress conditions influence its structure and propensity to aggregate (Martin et al., 2011, Oakley et al., 2020). Indeed, SFs and PHFs are formed from the pathological assembly of tau monomers to dimers, then oligomers and eventually fibrils, which deposit as NFTs. For example, while phosphorylation (Alonso et al., 1996), truncation (Wischik et al., 1988, Gamblin et al., 2003), or cysteine oxidation promote tau aggregation (Schweers et al., 1995), other modifications, such as nitration of tyrosine residues inhibit tau nucleation and/or elongation (Reynolds et al., 2005a, Reynolds et al., 2005b) and glycation stabilises filaments and increases insolubility of NFTs (Ko et al., 1999, Necula and Kuret, 2004).

The longest central nervous system isoform of human tau has five tyrosines, at positions 18, 29, 197, 310, and 394. In addition to nitration, these residues are targets of other post-translational modifications, including phosphorylation and dityrosine cross-linking. All of these alterations could result in the accumulation of oligomers (Lebouvier et al., 2009, Ait-Bouziad et al., 2020, Tremblay et al., 2010, Reynolds et al., 2005b, Maina et al., 2021). To understand the contribution of dityrosine formation on tau, we have recently reported a new tau model for *in vitro* studies of aggregation of a 95 amino acid tau fragment comprised of residues Ile-297–Glu-391 of the full-length tau, also referred to as dGAE (Al-Hilaly et al., 2017). The dGAE fragment was previously isolated from the proteolytically stable PHF core (Wischik et al., 1988) and overlaps with the core of AD PHFs revealed by cryo-electron microscopy (Fitzpatrick et al., 2017). dGAE assembles into filaments without the need for additives such as heparin or arachidonic acid (Al-Hilaly et al., 2017, Al-Hilaly et al., 2019a) and is able to form an identical structure to ex-vivo filaments extracted from AD and Chronic Traumatic Encephalopathy (CTE) (Lövestam et al., 2022). Using metal-catalysed oxidation (MCO), we previously showed that oxidation of dGAE (which contains tyrosine 310) results in rapid formation of dityrosine crosslinked tau oligomers, which lack the ability to elongate into fibrils, suggesting that the dityrosine bond formation results in the stabilisation of the oligomers (Maina et al., 2021), similar to Aβ oligomers (Maina et al., 2020). Consistent with this, ONOO^-^-treated full-length tau filaments in an arachidonic acid model of tau aggregation was previously shown to increase filament stability (Reynolds et al., 2006) similar to stability and insolubility of brain-derived PHFs (Kondo et al., 1988, Greenberg and Davies, 1990, Miao et al., 2019).

Given the high stability and irreversibility of the dityrosine bonding (Gross and Sizer, 1959), crosslinked proteins can be detected in various *in vitro* and *in vivo* settings. Aβ and α-synuclein have both been shown to form dityrosine cross-links in *vitro*, and dityrosine cross-linked Aβ and α-synuclein have each been localised in Aβ plaques in AD and in Lewy bodies in Parkinson’s disease (PD) post-mortem brain tissues, respectively (Galeazzi et al., 1999, Souza et al., 2000a, Al-Hilaly et al., 2013, Al-Hilaly et al., 2016). This raises the question of whether dityrosine cross-linking is present on human-derived tau assemblies. Using TEM immunogold labelling and fluorescence spectroscopy, we show that dityrosine cross-linking is observed on human AD tau oligomers and PHFs *in vivo*, and dityrosine colocalises with tau on NFTs in AD. However, using MCO of dGAE in vitro, we showed that prefibrillar tau assemblies are more amenable to dityrosine cross-linking than fibrils, suggesting that dityrosine crosslinking *in vivo* may have a different impact on the tau assemblies and activity. Nonetheless, dityrosine formation results in heat and SDS stability of oxidised prefibrillar and fibrillar tau assemblies, suggesting a general role of dityrosine crosslinking in enhancing pathological tau stability.

## Results

### Dityrosine crosslinking is observed in pathological tau from AD samples

We have previously shown the presence of dityrosine crosslinks in AD amyloid plaques (Al-Hilaly et al., 2013). Here, we first conducted immunogold labelling TEM on AD brain sections to investigate the presence of dityrosine crosslinks on pathological tau. The labelling of AD brain sections revealed densely packed tau containing filaments, many showing the characteristic paired-helical appearance upon close inspection (Figure 1A). These filaments also labelled with an anti-dityrosine antibody, indicating the presence of dityrosine crosslinks. Since PHFs were not found in the age-matched control brain sections, only the data from the AD patients is shown, although immunogold labelling confirmed no non-specific labelling was present for either antibody.

**Figure 1.**
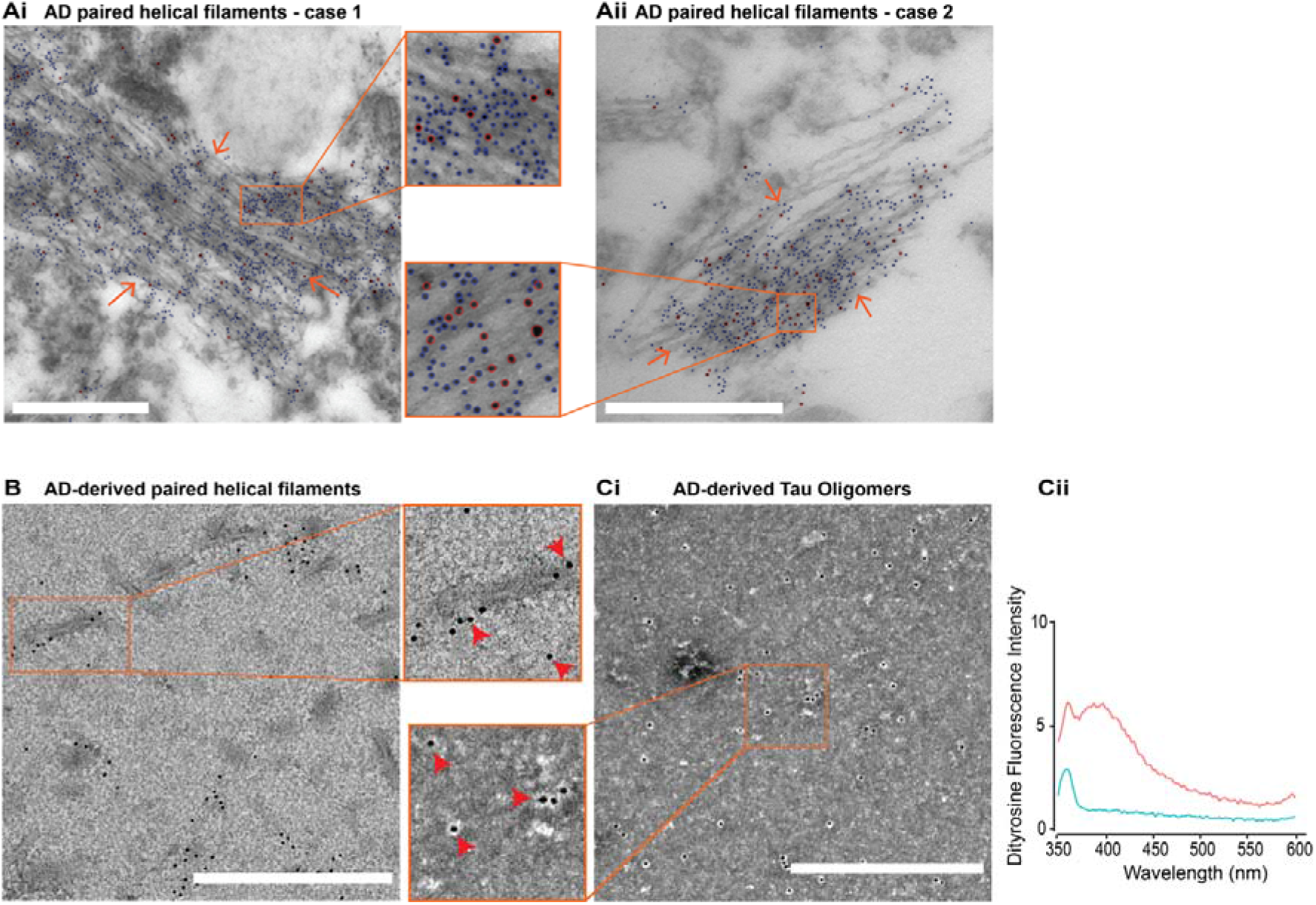
Dityrosine detection in AD-derived Tau species. **(A)** TEM double immunogold labelling revealed close, dense labelling of PHFs (orange arrow) using anti-dityrosine mouse monoclonal antibody (secondary gold conjugated anti-mouse 10 nm) and anti-tau rabbit polyclonal antibody (secondary gold conjugated anti-rabbit 5 nm) on AD brain sections of neurofibrillary tangles (Ai & Aii, see Inserts: blue circles for tau and red circles for dityrosine). Red arrows highlight PHF.. **(B)** TEM single immunogold labelling revealed labelling of anti-dityrosine mouse monoclonal antibody on ex-vivo AD-derived paired helical filaments (Insert: red arrows for dityrosine). **(Ci)** TEM single immunogold labelling revealed anti-dityrosine mouse monoclonal antibody labelling on ex-vivo AD-derived tau oligomers. (**Cii**) Dityrosine signal was confirmed for the AD-derived tau oligomers with fluorimeter using fluorescent excitation/emission 320nm/340 – 600 nm, with dityrosine peak signal observed between 400-420 nm. Scale bar = 500 nm.

Ex-vivo characterised AD tau filaments (Fitzpatrick et al., 2017) were also examined using immunogold TEM. Electron micrographs show the labelling of anti-dityrosine on the tau filaments supporting the view that the tau protein is cross-linked within the filaments (Figure 1B). To examine whether dityrosine crosslinks occur earlier in the assembly process, AD brain-derived oligomers (Lo Cascio et al., 2020, Lasagna-Reeves et al., 2012) were labelled with dityrosine antibody using single immunogold labelling, and these show the distinctive gold labels suggesting that dityrosine is also found within these earlier oligomeric species (Figure 1C). In support of the presence of dityrosine crosslinks within oligomeric tau, an emission peak was observed for the oligomeric species in solution at 410 nm, the expected signal for dityrosine (Figure 1D) (Atwood et al., 2004, Al-Hilaly et al., 2013, Maina et al., 2020). Together, these data reveal that dityrosine crosslinks appear in both tau oligomers and fibrils in vivo and ex-vivo, suggesting a potentially important role for dityrosine in tau assembly and neurofibrillary tangle accumulation in AD.

### Soluble and prefibrillar dGAE are more amenable to dityrosine crosslinking and its impact than fibrillar dGAE assemblies

To further examine the potential effects of dityrosine crosslinking on fibrillar tau, we examined the labelling of filaments formed from tau(297-391; also termed dGAE), which has been shown to form PHFs and SFs without further additives (Lövestam et al., 2022, Pollack et al., 2020, Al-Hilaly et al., 2019b, Lutter et al., 2022, Al-Hilaly et al., 2017). Agitation of 300 μM dGAE for 6 h at 37° C resulted in the formation of a mixture of oligomeric and pre-fibrillar species, while incubation for 48h resulted in mature filaments with a pronounced twist and some lateral bundling (Figure 2A, B, C).

**Figure 2.**
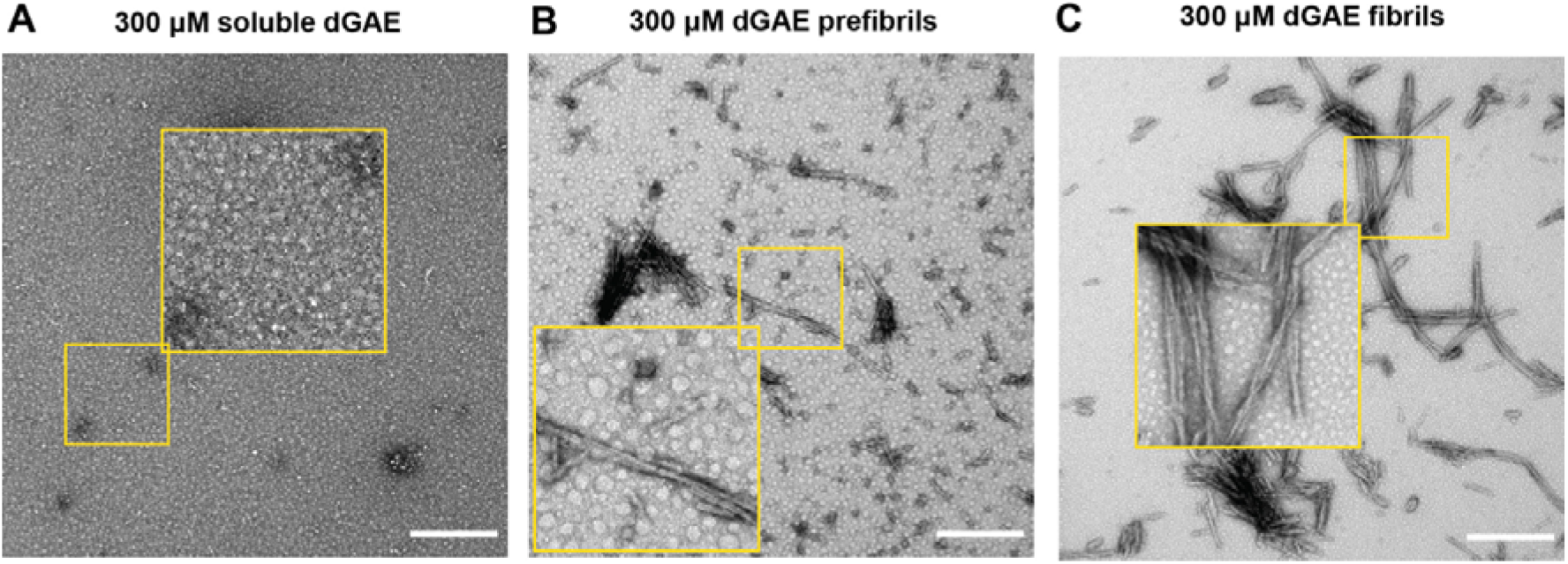
Preparation of dGAE assemblies for oxidation experiments. dGAE assemblies (300 μM) were prepared freshly (**A**), and incubated at 37°C with agitation at 700 RPM for 6 h **(B)** or 48 h **(C)**. TEM imaging for the freshly prepared sample revealed small, round assemblies, described here as soluble species (**A**). The samples incubated for 6 h revealed short fibrils and small round assemblies, described here as prefibrils (**B**). Samples incubated for 48 h revealed long twisted and bundled fibrils described here as fibrils (**C**). Scale bar = 500 nm.

We have shown that metal-catalysed oxidation (MCO) using Cu^2+^ and H_2_O_2_ results in dityrosine crosslinking of the dGAE (Maina et al., 2021). In this study, soluble, pre-fibrillar and fibrillar dGAE samples (50 μM) were prepared from their respective stock preparations (Fig. 2). The 50 μM soluble, pre-fibrillar and fibrillar forms of dGAE were incubated for 15 mins or 48h under MCO conditions using Cu^2+^ and H_2_O_2_ to induce the formation of dityrosine crosslinks. The tyrosine and dityrosine levels within the samples were evaluated together by fluorescence spectroscopy using an excitation wavelength of 280 nm for tyrosine and excitation of 310 nm for dityrosine (Figure 3A-D). Soluble dGAE shows a strong tyrosine fluorescence signal which is reduced substantially following a 15 min incubation in oxidising conditions (Fig. 3A), with the concurrent increase in the dityrosine signal at 410-420 nm (Fig. 3B). Similarly, prefibrils showed a transition to dityrosine fluorescence following oxidation (Fig. 3A-B). The level of dityrosine signal in the soluble and prefibrillar dGAE samples remained similar between 15 min and 48 h suggesting that the formation of dityrosine is rapid (Fig. 3B & D). In contrast, mature fibrillar dGAE showed an extremely low dityrosine signal at 15 min, which remained similar at 48h (Fig. B & D), suggesting that fibrils are less susceptible to dityrosine crosslinking. The tyrosine signal reduces from 15 min to 48h incubation for all samples which may be due to lateral association of fibrils with time leading to the burial of the tyrosine. Together, this reveals that soluble and prefibrillar dGAE assemblies are more amenable to dityrosine crosslinking than fibrillar species.

**Figure 3.**
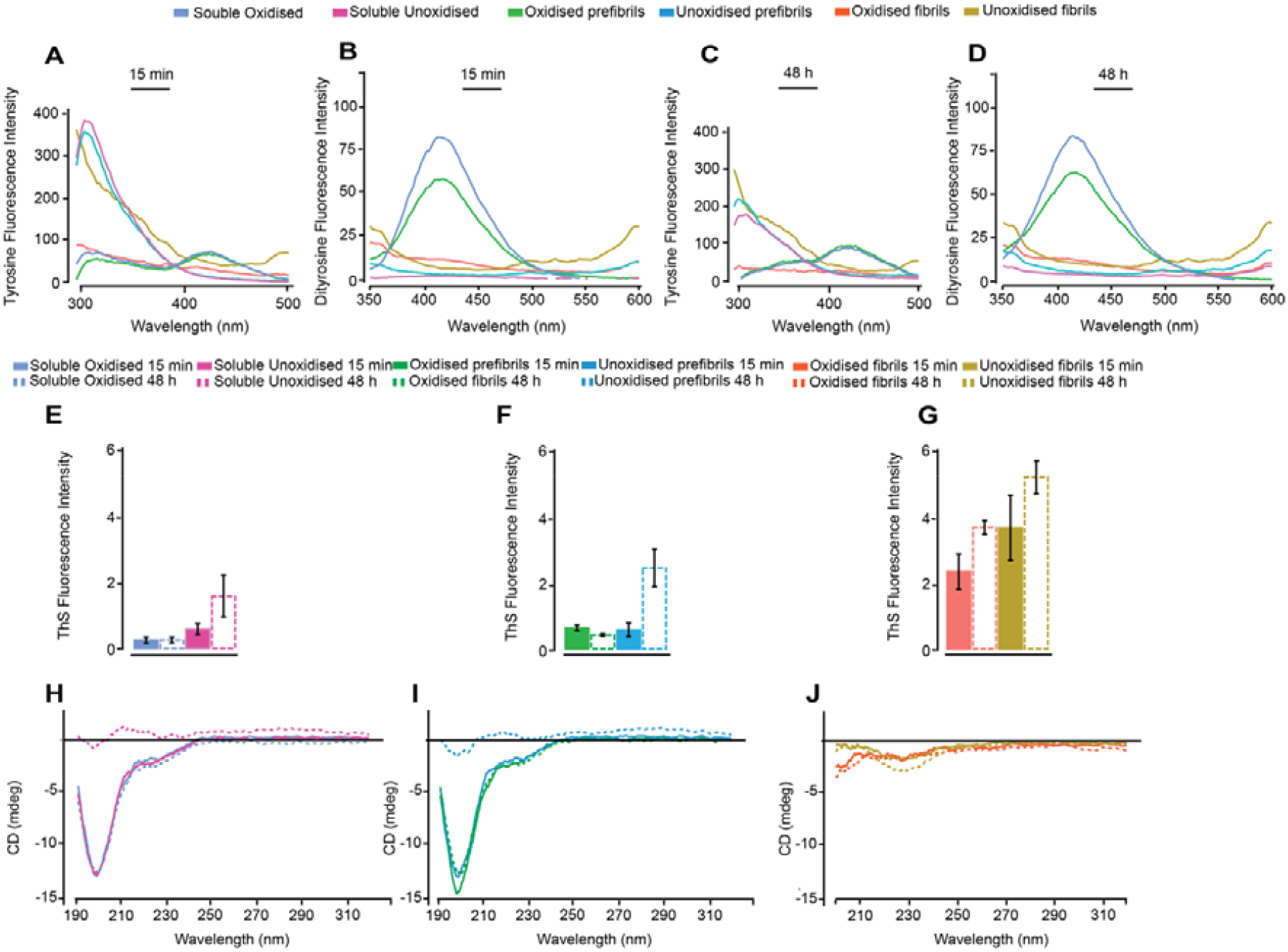
Self-assembly properties and conformation of oxidised dGAE assemblies. Soluble, prefibrils and fibrils of dGAE (50 μM) were prepared from their respective stock assemblies, left unoxidised or oxidised using CuCl_2_:dGAE at 10:1 ratio, then mixed with 2.5 mM H_2_O_2_ and left for 15 min or 48 h at 37°C/700 rpm. Tyrosine signal was monitored using an excitation/emission 280nm/290 – 500nm, with peak tyrosine signal observed at 305 nm. While dityrosine signal was collected using fluorescent excitation/emission 320nm/340 – 600 nm, with dityrosine peak signal observed between 400-420 nm. Tyrosine signal collected at 15 min incubation shows a decrease in intensity for all the oxidised samples (**A**). Fluorescence reading showed a high intensity signal for dityrosine formation in oxidised soluble assemblies, with lower intensity observed for prefibrils and fibrils (**B**). Tyrosine signal at 48 h showed a further decrease in all oxidised samples and a minor reduction in the unoxidised samples compared to signals collected at 15 min (**C**). Dityrosine signal increased further in comparison to 15 min signal for oxidised assemblies (**D**). ThS fluorescence assay was conducted to observe the degree of self-assembly using excitation/emission of 440/460-600. Soluble (**E**) prefibrillar (**F**) and fibrillar (**G**) unoxidised dGAE assemblies showed increased ThS fluorescence at 48 h compared to 15 min; while oxidised samples remained similar between the time points. Circular dichroism (CD) spectroscopy for all the soluble assemblies except soluble unoxidised samples at 48 h revealed a typical random coil signal (**H**). All dGAE prefibrils except the unoxidised prefibrils at 48 h, also showed random coil signal, however to a lower extent in unoxidised prefibrils at 15 min and oxidised prefibrils at 48 h (**I**). CD showed the absence of random coiled conformation in unoxidised and oxidised fibrils and the appearance of β-sheet, especially in unoxidised fibrils at 48 h indicated by the presence of a minimum at 228nm (**J**).

In order to examine the amyloid-fibrillar nature of the dGAE samples, ThS fluorescence assays were conducted. ThS fluorescence intensity increased only following 48 h incubation in the unoxidised soluble and prefibrillar dGAE assemblies Figure 3E-F) and no significant increase in the fluorescence intensity was observed in the oxidised samples. Both these observations are consistent with previous results whereby oxidation appears to halt further self-assembly (Maina et al., 2021). A minor increase in ThS fluorescence intensity was observed in the 48 h unoxidised fibrils compared to the same samples incubated for 15 min (Figure G). However, unlike the oxidised soluble and prefibrillar dGAE assemblies, oxidised dGAE fibrils showed increased ThS fluorescence intensity at 48 h when compared to the 15 min signal, suggesting that the fibrils still retain some ability to assemble further which may arise from their reduced susceptibility to crosslinking following oxidation.

To investigate the secondary structure of the assemblies, circular dichroism spectroscopy (CD) was conducted (Figure 3H-J). CD for unoxidized soluble and prefibrillar species appear to show random coil signal after 15 mins but progress to a lower solubility form after 48 h incubation as indicated by the low intensity signal observed. However, both soluble and prefibrillar oxidised samples show very strong random coil signal arising from soluble, random coil dGAE in solution which changes only slightly in intensity from 15 mins to 48 h incubation. CD spectra for fibril samples show the expected β-sheet signal for the unoxidized fibrils sample which increases with time, but the decreased intensity β-sheet signal for the oxidised sample may suggest lower solubility. This seems to suggest that changes happen rapidly and there is little further conformational change after 15 mins incubation following dityrosine crosslinking.

Soluble, prefibrillar and fibrillar samples of dGAE that had been oxidised for 15 mins only or remained unoxidised were compared using TEM. As expected, the electron micrographs of unoxidized dGAE samples showed very small particles for soluble dGAE, short fibrils for prefibrillar and longer fibrils for those categorised as fibrils (Figure 4 A-C). In contrast soluble (Figure A1 and A2) and prefibrillar (Figure B1 and B2) samples appeared to form small and larger rounded structures following oxidizing incubation, suggesting that that oxidation of amino acid residues including tyrosine to form dityrosine resulted in the prevention of further elongation and formation of trapped, relatively stable spherical species. However, the fibrillar sample remains fibrillar and their morphology changed very little compared to the unoxidized samples. Some lateral association of fibrils was apparent, and this may arise from crosslinking between fibres (Figure C1 and C2).

**Figure 4.**
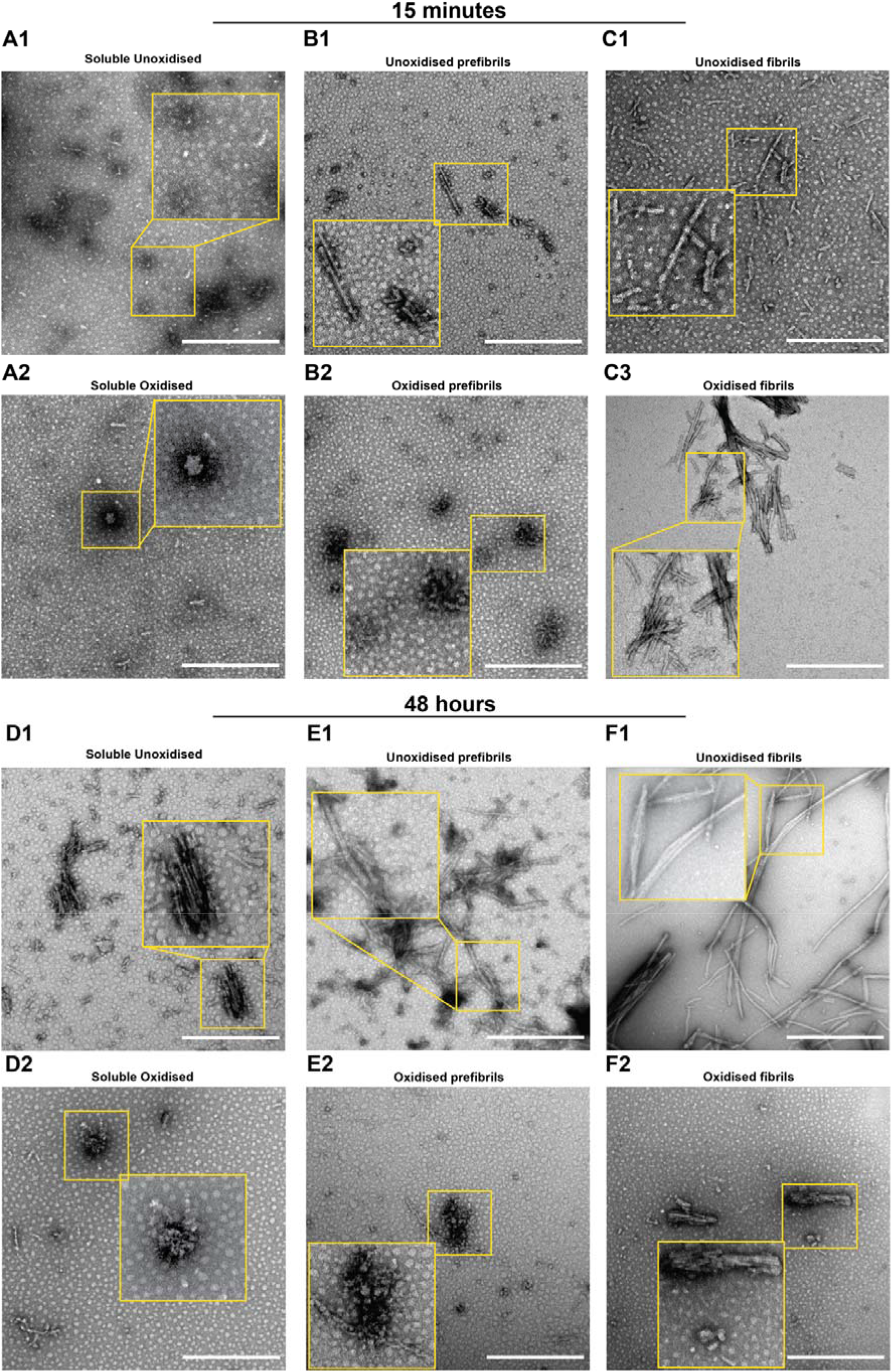
Morphologies of unoxidised and oxidised assemblies. Soluble, prefibrils and fibrils of dGAE (50 μM) were prepared from their respective stock assemblies, incubated unoxidized or oxidized using CuCl_2_:dGAE at 10:1 ratio, then mixed with 2.5 mM H_2_O_2_ and left for 15 min or 48 h at 37°C/700 RPM. Compared to the unoxidised soluble assemblies, oxidized soluble assemblies showed larger round assemblies at 15 mins (**A1-A2)**. Similarly, compared to the unoxidized prefibrils, oxidised prefibrils showed mostly large amorphous assemblies comprised of inter-connected smaller assemblies at 15 mins (**B1-B2**). Compared to unoxidised fibrils, oxidised fibrils showed many laterally connected fibrils at 15 min (**C1-C2**). The soluble unoxidised assemblies formed short fibrils at 48 h (**D1**), while the soluble oxidised assemblies formed large round assemblies (**D2)**. Similarly, unoxidised prefiibrils showed long, mature fibrils at 48 h (**E1**), while the oxidised prefibrils formed large round assemblies (**E2**). Unoxidised fibrils showed longer, matured fibrils (**F1**), while the oxidised fibrils revealed short, blunt morphology at 48 h (**F2**). Scale bar = 500 nm.

After 48h incubation, all three samples incubated under unoxidised conditions had assembled to some extent with either very long fibrils, or shorter fibres in the soluble dGAE sample (Figure 4 E-F). In comparison, after incubation in oxidising conditions, the soluble and prefibrillar samples showed mostly trapped, spherical species (similar to those found at 15mins incubation) but with some short fibrillar protofibrils (narrower elongated fibrils). The fibrillar sample showed shortening of the fibrils, suggestive of fragmentation or breakage of the stabilised fibres, or prevention of elongation, with more obvious lateral association than observed after 15 mins incubation. This may indicate that these cross-linked stabilised fibres are susceptible to breakage during incubation with shaking.

To investigate the apparent stability caused by the dityrosine crosslinking further, we investigated the heat and SDS insolubility of the oxidised and unoxidised samples (Figure 5A). The samples were boiled at 100 C in Laemmli sample buffer containing 4.4% LDS and 10% β-mercaptoethanol and then resolved by SDS−PAGE electrophoresis and stained with Coomassie brilliant blue (Reynolds et al., 2006). Unoxidised soluble dGAE sample mainly separated as a monomer (10/12 kDa) while prefibrillar and fibrillar samples show the appearance of a dimer (20 kDa/24 kDa). Insoluble species are found in the well and are most clearly evident in the fibril sample as expected. For the oxidized samples, there is a clear increase in the protein material in the well for each soluble, prefibrillar and fibril sample compared to the unoxidized and a dimer appears in all three samples. The fibril sample also shows some higher molecular weight species which suggests the stabilization of larger species potentially via dityrosine.

**Figure 5.**
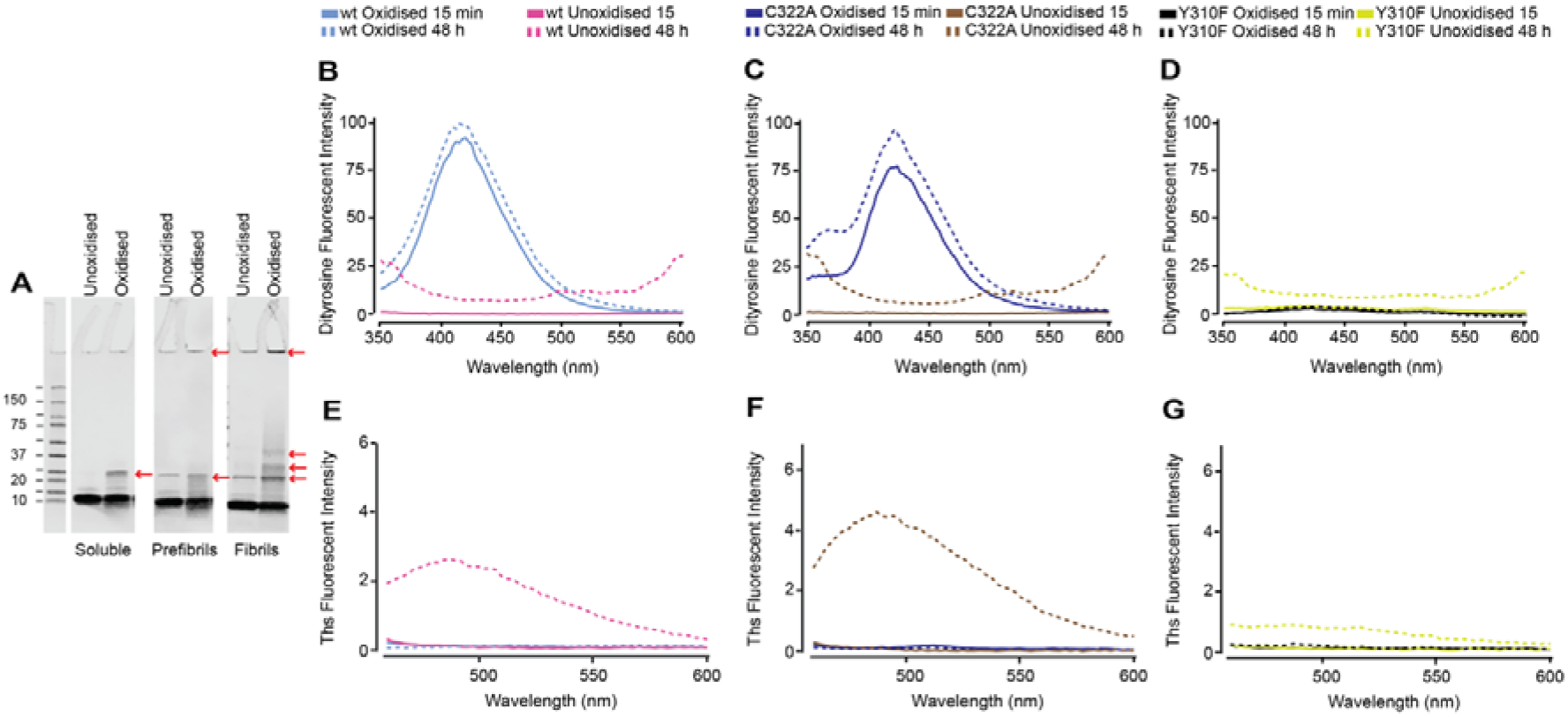
Stability of crosslinked assemblies, dityrosine formation susceptibility and self-assembly in dGAE variants. Soluble, prefibrils and fibrils of dGAE (50 μM) were prepared from their respective stock assemblies, left unoxidised or oxidised using CuCl_2_:dGAE at 10:1 ratio, then mixed with 2.5 mM H_2_O_2_ and left for 15 min at 37°C/700 rpm. Compared to all unoxidised assemblies, the oxidised assemblies showed heat and SDS insolubility that is more pronounced in the fibril sample (**A)**. 50 μM soluble wild-type dGAE, dGAE with C322A substitution and dGAE with Y310F substitution were prepared, left unoxidised or oxidised using CuCl_2_:dGAE at 10:1 ratio then mixed with 2.5 mM H_2_O_2_ and left for 15 min or 48h at 37°C/700 rpm. Fluorescence reading showed an increase in intensity signal for dityrosine in oxidised wildtype dGAE (**B**) and C322A dGAE (**C**) samples at 15 min and 48 h. No dityrosine signal was detected in Y310F dGAE sample (**D**). Unoxidised wildtype dGAE (**E**) and C322A dGAE (**F**) assemblies incubated for 48 h showed increased ThS fluorescence suggesting assembly, while Y310F dGAE showed no self-assembly (**G**). In contrast, oxidised samples showed no ThS fluorescence suggesting prevention of assembly.

This shows that 15 mins oxidation alone results in heat and SDS insolubility of the soluble, prefibrillar and fibrillar assemblies. This is more pronounced in the fibrillar sample, which indicated that heating and incubation in the Laemmli buffer resulted in the dissociation of the majority of the unoxidised fibrils to monomers and multimers. However, the oxidised samples that were treated the same way showed assemblies that remained as monomers, multimers, and some were too large and would have failed to enter the gel (≥250 kD, red arrows). This suggests that the oxidised assemblies are stabilised through dityrosine formation.

dGAE contains a single cysteine residue at residue 322. To examine whether the oxidation of cysteine plays a role in the conformational and morphological changes in dGAE upon oxidation, the ability of C322A variant dGAE to form dityrosine was investigated and compared with a control dGAE with a Y310A substitution. Oxidised wildtype dGAE shows a strong signal from dityrosine at 410-420 nm as previously shown, and this remains clearly observed for C322A dGAE with a similar intensity as wildtype (Figure 5 B-C). As expected, however, this signal is not observed for Y310F since this variant does not include a tyrosine residue. ThS fluorescence shows that wildtype dGAE and C322A dGAE assemble over 48 h only when incubated under non-oxidizing conditions, while Y310F appears to be unable to self-assemble under either oxidising or non-oxidising conditions under conditions used in this study (Figure 5 E-G). Collectively these data showed that cysteine is not involved in the dGAE stabilisation following oxidation, while tyrosine appears to be involved in dGAE assembly.

## Discussion

Dityrosine occurs naturally in proteins, helping to increase their stability. This modification has been found in many elastic and structural proteins, including elastin, fibroin, keratin, cuticlin, and collagen (Labella et al., 1967, Raven et al., 1971, Fujimoto, 1975, Waykole and Heidemann, 1976), where it is believed to increase their mechanical strength and subsequently their insolubility (Skaff et al., 2005). However, the formation of dityrosine in proteins is also implicated in many diseases, including AD, and PD (Souza et al., 2000b, Atwood et al., 2004, Al-Hilaly et al., 2013, Al-Hilaly et al., 2016), cystic fibrosis (Van Der Vliet et al., 2000), atherosclerosis (Leeuwenburgh et al., 1996), cataracts in the eye lens (Wells-Knecht et al., 1993, Bodaness and Zigler, 1983) and acute myocardial infarction (Mayer et al., 2014). Our previous work has co-localised dityrosine with Aβ within Aβ plaques (Al-Hilaly et al., 2013) and Lewy bodies with α-synuclein (Souza et al., 2000a, Al-Hilaly et al., 2016), suggesting that it could play a role in increasing the insolubility of Aβ plaques in AD and α-synuclein aggregates in PD. It remained to be determined whether dityrosine crosslinking is formed in the tau aggregates in human AD brains. Here, we have revealed that dityrosine forms *in vivo* on tau oligomers and filaments within NFTs both in situ in brain and also localised to ex-vivo AD-derived assemblies. Using the PHF-core tau fragment (dGAE) as a model, which forms dityrosine cross-links via tyrosine 310, we also showed that although dityrosine forms on tau assemblies, mature fibrillar species are less amenable to dityrosine crosslinking and that dityrosine formation promotes heat and SDS insolubility of the tau aggregates.

It is known that exposure to reactive oxygen species (ROS), ageing, nitrogen dioxide, metal ions and lipid hydroperoxides can result in dityrosine formation (Giulivi and Davies, 1993, Giulivi and Davies, 1994, Kato et al., 1994, Atwood et al., 2004). Oxidative stress accumulates early in AD, increasing with disease pathology (Nunomura et al., 2001, Butterfield and Halliwell, 2019). ROS, ageing, metal ions and other factors could induce dityrosine crosslinking on the tau oligomers and fibrils *in vivo*. Dityrosine crosslinking on tau fibrils could increase their stability and insolubility. The PHFs extracted from the AD brain are highly insoluble and resistant to proteolytic cleavage. Specifically, the early-stage PHF-tau has reduced SDS solubility (Lee et al., 1991, Greenberg and Davies, 1990), while late-stage PHF tau exhibits SDS and sarcosyl insolubility (Kondo et al., 1988, Greenberg and Davies, 1990, Miao et al., 2019). Previous work on full-length tau aggregation with arachidonic acid showed that dityrosine crosslinked tau fibrils have increased SDS stability, suggesting that the crosslinking may play a role in stabilising the early-stage PHFs and confers stability to convert into late-stage PHFs (Reynolds et al., 2006). Here, we find that the oxidised dGAE fibrils are difficult to resuspend in solution, are heat and SDS-insoluble and remained strongly laterally associated to one another, shown by TEM. Given that we find dityrosine crosslinking on AD-derived tau fibrils and in NFTs, this finding suggests that this post-translational modification increases the insolubility of tau fibrils and NFTs in AD.

By comparing the capability of soluble, prefibrillar and fibrillar dGAE to form dityrosine, our findings indicate that fibrillar tau assemblies are less amenable to dityrosine crosslinking. This can be seen by the very low fluorescence signal observed in the oxidised fibrillar dGAE samples in spectroscopic experiments, unlike in the oxidised soluble and prefibrillar assemblies, which showed robust dityrosine formation. This apparent inability to form dityrosine compared to soluble and prefibrillar assemblies could be due to the inaccessibility to the tyrosine residue on the tau molecule when it is arranged in a fibrillar architecture. The only tyrosine present in dGAE is at position 310, which is buried within the PHF core in the Cryo-EM structures of tau filaments from many tauopathies, including AD and CTE (Ait-Bouziad et al., 2020)(Fitzpatrick et al., 2017, Zhang et al., 2019, Shi et al., 2021, Lövestam et al., 2022). Recent Cryo-EM evidence indicates that the dGAE forms a similar structure as that of AD and CTE (Lövestam et al., 2022). Therefore, it can be concluded that the decreased ability of dGAE fibrils to form dityrosine under MCO conditions is due to the inaccessibility of the tyrosine residue, unlike in the soluble and prefibrillar dGAE assemblies.

Our *in vitro* work on dGAE and Aβ both suggest that dityrosine formation traps the assemblies in a conformation that doesn’t favour elongation into fibrils (Maina et al., 2020, Maina et al., 2021). However, whether dityrosine crosslinked tau oligomers also fail to elongate to fibrils *in vivo* is a question of future research. We have shown that dityrosine crosslinked tau oligomers are not toxic to human neuroblastoma cells even after three days in culture (Maina et al., 2021) although we also find that dGAE is generally non-toxic in its oligomeric form (Pollack et al., 2020). It is unclear at the moment what the impact of dityrosine crosslinking is on the aggregation and toxicity of tau oligomers *in vivo*. Nonetheless, it is essential to highlight that the influence of dityrosine on tau aggregation and toxicity may depend on the extent of the dityrosine crosslinking (Maina et al., 2021). A highly crosslinked protein may be highly trapped and exhibit a different property than a partially crosslinked protein. This, therefore, makes it impossible to speculate a general model of the influence of this modification on tau and indeed other proteins like Aβ.

Finally, cysteine oxidation can be induced by oxidative stress conditions and is known to influence the functions, properties and toxicity of tau (Landino et al., 2004, Saito et al., 2021, Weismiller et al., 2021). dGAE has a single cysteine residue at position 322, and we have previously shown that it forms PHFs independent of this cysteine and that assembly is enhanced under reducing conditions or using C322A variant (Al-Hilaly et al., 2017, Al-Hilaly et al., 2018). To examine whether the oxidation of cysteine plays a role in the conformational and morphological changes in dGAE upon oxidation, we utilised dGAE C322A. This revealed that cysteine oxidation is not involved in stabilising the tau aggregates; instead, the effects observed resulted from dityrosine crosslinking. However, this does not rule out the involvement of other oxidative modifications, such as the oxidation of histidine and phenylalanine, which are all present in dGAE. Future studies would delineate the specific amino acids involved. Nonetheless, the immunogold labelling experiments on AD brain tissue and AD-derived fibrils and oligomers strongly suggest that dityrosine crosslinking has *in vivo* relevance to tau in AD, similar to its role on Aβ (Al-Hilaly et al., 2013) and α-synuclein (Souza et al., 2000a, Al-Hilaly et al., 2016) in AD and PD, respectively. Given the role of dityrosine in increasing the mechanical strength and insolubility of proteins (Skaff et al., 2005), we believe that its association with hallmarks of AD (tau and Aβ) and PD (α-synuclein) that are known to be insoluble in these diseases, could suggest its participation in promoting the insolubility, accumulation and resistance to degradation of NFTs and Aβ plaques in AD and alpha-synuclein aggregates in Lewy bodies in PD.

In conclusion, for the first time, our findings showed that, similar to Aβ plaques and α-synuclein Lewy bodies, dityrosine crosslinks PHFs in AD and is a posttranslational modification of AD-derived tau oligomers and fibrils. Our finding suggests that the assembly state of the tau molecule will influence its ability to be crosslinked, such that, when the tyrosine is buried in the case of fibrils, it becomes less amenable to dityrosine formation. This finding has implications for understanding the mechanism governing the insolubility and toxicity of tau assemblies *in vivo* and providing therapeutic avenues that focus on the tau molecule.

## Materials and Methods

### Cases

Tissue from the middle---frontal gyrus of three AD subjects and one age-matched control were used for transmission electron microscope (TEM) immunogold labelling. The samples were obtained from the London Neurodegenerative Disease Brain Bank with informed consent and under material transfer agreement and stored at −80°C under local ethics committee guidelines (Table 1).

**Table 1.**
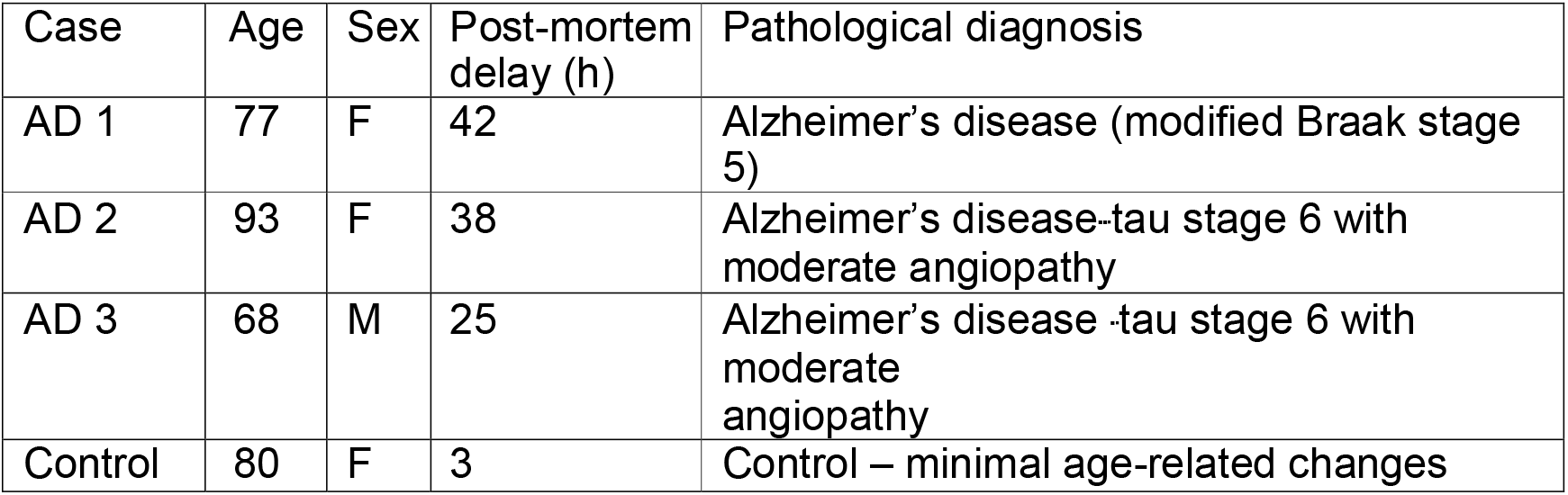
Characteristics of subjects.

| Case | Age | Sex | Post-mortem delay (h) | Pathological diagnosis |
| --- | --- | --- | --- | --- |
| AD 1 | 77 | F | 42 | Alzheimer's disease (modified Braak stage 5) |
| AD 2 | 93 | F | 38 | Alzheimer's disease-tau stage 6 with moderate angiopathy |
| AD 3 | 68 | M | 25 | Alzheimer's disease -tau stage 6 with moderate angiopathy |
| Control | 80 | F | 3 | Control – minimal age-related changes |

### Immunogold labelling for total Tau and Dityrosine

The brain sections were labelled using our established method of immunogold labelling (Al-Hilaly et al., 2013). Phosphate-buffered saline (pH 8.2) containing 1% BSA, 500 μl/l Tween-20, 10 mM Na EDTA, and 0.2 g/l NaN3 (henceforth called PBS+), was used throughout. Briefly, thin brain sections were mounted on TEM grids and blocked for 30 minutes in normal goat serum (1:10 dilution), labelled with 10 μg/ml IgG anti-tau rabbit polyclonal antibody (Sigma, SAB4501831) and 10 μg/ml IgG anti-dityrosine mouse monoclonal antibody (JaICA, Shizuoka, Japan), and left in a humid container overnight at room temperature. The sections were rinsed in PBS+ three times for 2 minutes and immunogold labelled with goat anti-mouse 10nm and goat anti-rabbit 5nm for 1 hour (1:10 dilution). Then the grids were rinsed three times for 10 minutes with PBS+, followed by four 5-minute rinses in distilled water, before post-staining in 0.05% aqueous uranyl acetate for 1 hour. Ex-vivo PHFs and tau oligomers were provided by Michel Goedert and Rayez Kayed groups, respectively. These samples have been well-characterised by both groups (Fitzpatrick et al., 2017, Lo Cascio et al., 2020, Lasagna-Reeves et al., 2012).

For the single labelling of ex-vivo PHFs and tau oligomers with dityrosine antibody, 4 μL of each sample were placed onto 400-mesh carbon-coated grids (Agar Scientific, Essex, UK), allowed to adhere for 1 min, and the excess sample removed using filter paper. The grids were blocked using normal goat serum (1:10 in PBS+) for 15 min, then incubated with mouse anti-dityrosine monoclonal antibody (10 μg/ml IgG; JaICA, Shizuoka, Japan) for 2 h at room temperature. The grids were rinsed three times for 2 min in PBS+, and then labelled with a 10-nm gold particle-conjugated goat anti-mouse IgG secondary probe (GaM10 British BioCell International, Cardiff, UK; 1:10 dilution) for 1 h at room temperature. The grids were rinsed five times for 2 min using PBS+, five times for 2 min with distilled water, then negatively stained as described below.

### Negative-stain transmission electron microscopy (TEM)

Samples (4 μL) were placed on 400-mesh carbon-coated grids (Agar Scientific, Essex, UK) and incubated for 1 min incubation. The excess sample was blotted using filter paper, and the grids were washed with 4 μL filtered milli-Q water. The grids were then negatively stained for 40 sec using 4 μL of filtered 2% (w/v) uranyl acetate. The excess stain was blotted with filter paper and grids left to air-dry before storage. The grids were examined on a JEM-400-plus transmission electron microscope (Jeol, USA), operated at 80 kV fitted with a Gatan Orius SC100 camera (UK).

### Preparation and assembly of dGAE

Recombinant wild-type and variant dGAE were prepared using our established method (Al-Hilaly et al., 2017). The dGAE was expressed in *Escherichia coli* and purified by P11 phosphocellulose chromatography following heat treatment. The protein fractions were eluted with 50 mM PIPES buffer (pH 6.8) or 50 mM MES buffer (pH 6.25), both supplemented with 1 mM EGTA, 5 mM EDTA, 0.2 mM MgCl_2_ and 5 mM 2-mercaptoethanol containing 0.1–1 M KCl. The peak of protein elution was identified by protein assay (at 0.3–0.5 M KCl) and dialyzed against 80 mM PIPES buffer (pH 6.8), 1 mM EGTA, 5 mM EDTA, 0.2 mM MgCl_2_, 5 mM 2-mercaptoethanol, or PB (10 mM; pH 7.4). The dGAE protein concentration was measured using Advanced Protein Assay Reagent (Cytoskeleton, Inc.) with bovine serum albumin as a standard. dGAE was finally diluted with 10 mM phosphate buffer (pH 7.4). dGAE (300 μM) was prepared in phosphate buffer (10mM; pH7.4) and incubated in the dark for i) 0 minutes; ii) 6 hours; or iii) 48 hours at 37°C with shaking at 700 rpm using a thermomixer C (Eppendorf, Germany). Based on TEM observations, henceforth, the samples will be referred to as (i) soluble dGAE; (ii) prefibrillar dGAE; and (iii) fibrillar dGAE, respectively.

### Metal-catalysed oxidation of dGAE

The soluble, prefibrillar and fibrillar dGAE samples were diluted to 50 μM before being used for metal-catalysed oxidation. For the fibrillar samples, 50 μM of the fibrils were obtained by first determining the concentration of the fibrils. This was achieved by spinning the fibrillar dGAE for 30 minutes at 20,000 g to separate the supernatant (containing unaggregated, soluble dGAE) and pellet (containing fibrils). The concentration of the soluble dGAE in the supernatant was used to back-calculate the concentration of dGAE in the fibrils. Soluble, prefibrillar and fibrillar dGAE (at 50 μM) were incubated in phosphate buffer (10 mM; pH7.4) alone (unoxidised) or in the presence of 500 μM CuCl_2_ (peptide: CuCl_2_ ratio 1:10) and 2.5 mM H_2_O_2_ (oxidised) and incubated for 15 minutes or 48 hours at 37°C with shaking at 700 rpm. The oxidation reaction was quenched by adding 2 mM EDTA to the assembly mixture. Oxidation of variant dGAE (carrying the mutations C322A and Y310F) was done using the same procedure. A minimum of three independent experiments were conducted to ensure the reproducibility of the findings.

### Fluorescence spectroscopy

Dityrosine formation was monitored using a fluorescent excitation wavelength of 320 and emission collected between 340 – 600 nm, with dityrosine peak signal expected between 400-420 nm, using a fluorescence spectrophotometer (Varian Ltd., Oxford, UK) and a 1-cm path length quartz cuvette (Starna, Essex, UK). The tyrosine fluorescence signal was monitored using an excitation wavelength of 280 nm and emission between 290 – 600, with the peak tyrosine emission observed at 305 nm. For all the measurements, the excitation and emission slits were set to 10 nm with a scan rate set to 300 nm/min with 2.5 nm data intervals and an averaging time of 0.5 s. The photomultiplier tube detector voltage was set at 500 V.

### Thioflavin S (ThS) fluorescence assay

To assess the assembly of dGAE in the different samples, a portion (10 μL) of each was incubated for 3 min with ThS (5 μM in 20 mM MOPs buffer, pH 6.8). ThS fluorescence intensity was observed using a fluorescence spectrophotometer (Varian Ltd., Oxford, UK) and a 1-cm path length quartz cuvette (Starna, Essex, UK). The excitation wavelength was set at 440 nm, with emission between 460 to 600 nm collected, with the peak emission observed at 483 nm. Each experiment included a minimum of three independent experiments. The peak emission was collected and analysed, then plotted graphically.

### Circular Dichroism (CD)

The secondary structure of the samples was assessed using the Jasco J715 CD spectrometer (Jasco, Goh-Umstadt, Germany). Each sample (40 μL) was placed into a 0.2-mm path length quartz cuvette (Hellma) and scanned between 190 and 260 nm. The CD spectra were collected in triplicate at a temperature of 21 °C.

### SDS-PAGE and immunoblotting

dGAE samples (20 μL) for each condition were mixed with 4x Laemmli sample buffer (containing 4.4% LDS) (Bio-Rad), supplemented with 10% 2-mercaptoethanol, boiled for 10 min at 100°C, resolved using a 4%–20% gradient Mini-Protean® TGX Precast Gels (Bio-Rad) at 100 V until the sample buffer reached the end of the gel. The gel was stained using Coomassie Blue stain. Images were collected using LI-COR Odyssey XF.

## Author contributions

M.B.M. planned and carried out the work; Y.K.A.-H., SO, TK, LB, KM and J.E.R. contributed experimental work. M.B.M. and L.C.S. wrote the paper; Y.K.A.-H and G.B. edited the paper and provided training. GB, CRH and CMW reviewed and edited the paper. LCS managed the project.

## Acknowledgements

The authors are grateful to M Goedert (MRC Laboratory of Molecular Biology, Cambridge, UK) for ex-vivo paired helical filaments which were kindly contributed by Bernandino Ghetti (Indiana University, USA). The authors are very grateful to Urmi Sengupta, Nemil Bhatt and Rakez Kayed (University of Texas, USA) for providing ex-vivo tau oligomers. TEM work was performed at the University of Sussex’s Electron microscopy imaging centre (EMC), funded by the School of Life Sciences, the Wellcome Trust (095605/Z/11/A, 208348/Z/17/Z) and the RM Phillips Trust. The authors thank Dr Pascale Schellenberger for valuable support.

## Data Sharing

No big data is generated from this work. The data generated and used to make figures in this manuscript will be available upon request.

